# Two types of human TCR differentially regulate reactivity to self and non-self antigens

**DOI:** 10.1101/2022.04.27.489747

**Authors:** Assya Trofimov, Philippe Brouillard, Jean-David Larouche, Jonathan Séguin, Jean-Philippe Laverdure, Ann Brasey, Gregory Ehx, Denis-Claude Roy, Lambert Busque, Silvy Lachance, Sébastien Lemieux, Claude Perreault

## Abstract

Based on analyses of TCR sequences from over 1,000 individuals, we report that the TCR repertoire is composed of two ontogenically and functionally distinct types of TCRs. Their production is regulated by variations in thymic output and terminal deoxynucleotidyl transferase (TDT) activity. Neonatal TCRs derived from TDT-negative progenitors persist throughout life, are highly shared among subjects, and are polyreactive to self and microbial antigens. Thus, >50% of cord blood TCRs are responsive to SARS-CoV2 and other common pathogens. TDT-dependent TCRs present distinct structural features and are less shared among subjects. TDT-dependent TCRs are produced in maximal numbers during infancy when thymic output and TDT activity reach a summit, are more abundant in subjects with AIRE mutations, and seem to play a dominant role in graft-versus-host disease. Factors decreasing thymic output (age, male sex) negatively impact TCR diversity. Males compensate for their lower repertoire diversity via hyperexpansion of selected TCR clonotypes.

## INTRODUCTION

Jawed vertebrates absolutely need a diversified TCR repertoire because classic αβ T cells must respond with exquisite specificity to an enormous diversity of ligands (Mittelbrunn and Kroemer, 2021). TCR diversity is generated by somatic recombination of V(D)J gene segments and is further increased postnatally by nucleotide insertion mediated by terminal deoxynucleotidyl transferase (TDT). Notably, neonatal thymocytes, which derive from fetal hematopoietic stem cells, do not express TDT (Rudd, 2020). TDT expression reaches maximal expression in humans between 10 and 40 months (mo) of age, then decreases progressively during adolescence and adulthood (Deibel et al., 1983; Pahwa et al., 1981). Recent estimates of the potential number of TCRs produced by V(D)J recombination range from 10^15^ (Mayer et al., 2019) to 10^61^ (de Greef et al., 2020), which vastly outnumbers the number of distinct TCRs present in a human body. Indeed, the adult human body contains approximately 4 × 10^11^ T cells (Jenkins et al., 2010) composed of about 10^10^ TCR clonotypes of various sizes (de Greef et al., 2020; Lythe et al., 2016). Initially, T cell repertoires have been presumed to be almost entirely private, and the occurrence of the same TCR in two unrelated individuals was attributed to coincidence. However, with the development of high-throughput TCR sequencing and state-of-the-art analytical algorithms, it became clear that interindividual sharing of TCR clonotypes was more common than expected (Pogorelyy et al., 2017; Sethna et al., 2019; Soto et al., 2020). Furthermore, some public clones were found to persist through an individual’s life (Chu et al., 2019; Pogorelyy et al., 2017). Still, the extent of interindividual sharing of TCR clonotypes is not precisely known (Johnson et al., 2021).

Which factors influence TCR diversity? At face value, the reduced thymic output associated with aging and male sex (Clave et al., 2018) should impinge on TCR diversity.

However, in contrast to mice, humans can compensate for a reduction of thymic output via minimal adjustments in homeostatic T cell proliferation (Goronzy and Weyand, 2019). Hence, the relation between thymic output and TCR diversity may not be linear. Nonetheless, analyses of TCR sequences in large cohorts have revealed a negative impact of aging on TCR diversity, while the effect of sex remains questionable (DeWitt et al., 2018; Krishna et al., 2020). Furthermore, there is an agreement that HLA polymorphism (i.e., heterozygosity for divergent alleles) positively correlates with TCR diversity and that some pathogens (e.g., CMV) can influence the composition of the TCR repertoire (DeWitt et al., 2018; Krishna et al., 2020).

In this study, we analyzed the physical properties of CDR3 beta sequences in seven cohorts of individuals and their implication in immune responses against pathogens, autoantigens, and alloantigens. We found stark differences between male and female TCR repertoires, where males maintain a lower diversity but high clonality (i.e., highly abundant TCR clonotypes). In comparison, female repertoires have a higher diversity of CDR3 at low clonal frequencies. A salient finding was the identification of two non-redundant CDR3 repertoire layers based on physical characteristics, including length, number of insertions, and V/J gene usage. The neonatal layer constitutes the entire TCR repertoire of cord blood, while the TDT-dependent layer appears later in life. Unexpectedly, the cord blood TCR repertoire contains mainly public and polyreactive CDR3s that are responsive to common pathogens.

## RESULTS

### Physical characteristics of public and superpublic CDR3s

We defined as *public* a CDR3aa (CDR3 amino acid sequence) seen in at least two individuals, while a *superpublic* CDR3aa is present in at least half of the subjects. For our first experiment, we used the Britanova cohort (Figure 1A), consisting of 79 healthy volunteers aged from 0-100 (see Methods). We found 2,862,268 public and 15,088 superpublic CDR3aa, of which 21 were ubiquitous (present in all samples) (Figure 1A). To define the physical properties of public and superpublic CDR3aa, we first analyzed their V and J gene usage by grouping the CDR3aa sequences by the annotated V or J gene identity. As expected (De Simone et al., 2018), while each unique CDR3aa sequence was encoded by mostly 1 or 2 J genes, many V genes can contribute to the same CDR3aa sequence. At the population level, we observed an average of 26 different V genes per public CDR3aa sequence (Figure 1B,C). For both public and superpublic CDR3aas, sequences encoded by a higher diversity of J genes were also encoded by numerous V genes (Figure 1D,E). In single individuals, up to eight different V genes could contribute to the same CDR3aa (Figure 1G). Finally, as previously reported (Gil et al., 2020; Madi et al., 2014; Venturi et al., 2008), we confirmed a positive correlation between the extent of CDR3aa sharing and the number of different nucleotide sequences encoding each CDR3aa (Figure 1F). This positive correlation points towards a trend of convergent recombination for public and superpublic CDR3aa (Quigley et al., 2010). We then used the software IgBLAST to obtain the number of mismatched (i.e., not germline) nucleotides in each CDR3aa sequence in the Britanova cohort (see Methods). We found that sequences shared by more individuals were also sequences with fewer mismatches (Figure 1H). This is consistent with the idea that non-templated nucleotide addition is a random process, and therefore each nucleotide mismatch lowers the likelihood of sequence sharing (Marcou et al., 2018; Sethna et al., 2019). Using the OLGA software (see Methods), we calculated the probability of a CDR3 nucleotide sequence being generated during V(D)J recombination for individual CDR3aa sequences. We confirmed a positive correlation between sequence publicness and recombination probability (Figure 1I). Finally, public sequences were shorter than private ones (Figure 1J), presumably because non-templated nucleotide addition lengthens the sequence (Marcou et al., 2018; Sethna et al., 2019).

**Figure 1:**
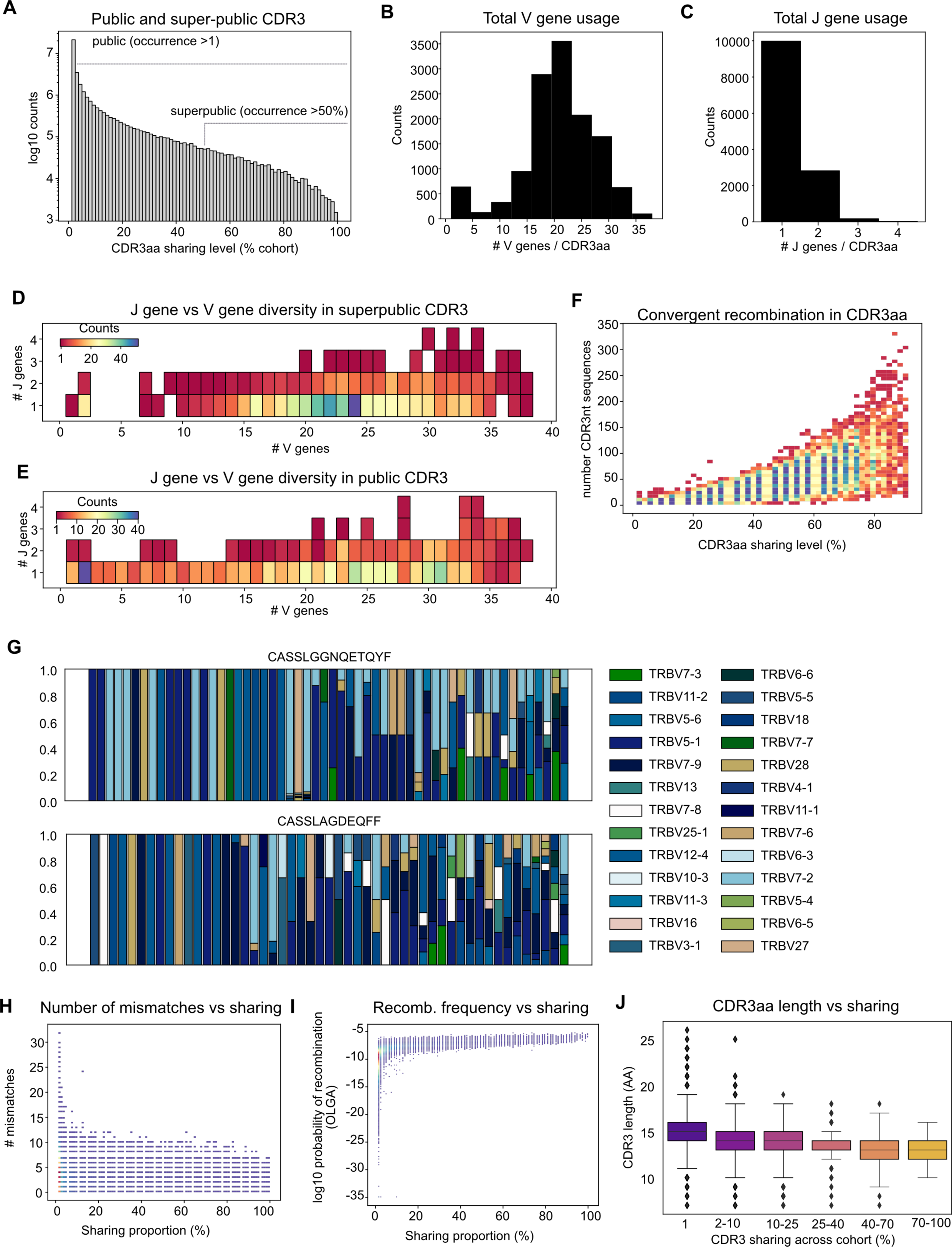
The physical characteristics of public CDR3s. **(A)** Public CDR3s are defined as seen in at least two people in the cohort, while superpublic CDR3s are seen in at least half of the cohort. Number of unique **(B)** V and **(C)** J genes encoding individual public CDR3aa. Relationship between the number of unique V and J genes shown by 2D histogram in **(D)** superpublic and **(E)** public CDR3aa. **(F)** Histplot showing convergent recombination of CDR3 nucleotide sequences in public CDR3aa: highly shared CDR3aa are coded by multiple synonymous nucleotides sequences. **(G)** Intra-individual V gene usage diversity for two superpublic CDR3aa sequences, sorted by intra-individual entropy: CASSLAGDEQFF and CASSLGGNQETQYF. **(H)** CDR3aa sharing and number of mismatches to the annotated germline. **(I)** Relation between the predicted recombination frequency and CDR3aa cohort sharing percentage. **(J)** CDR3aa length binned by CDR3aa cohort sharing percentage.

### CDR3aa sharing patterns change with age

Analysis of the Britanova cohort revealed a tight correlation between the extent of CDR3aa sharing among subjects and the frequency of the corresponding clonotypes in individual subjects. Superpublic CDR3aa were coded by high-frequency TCR clonotypes, and the most superpublic CDR3aa were found at higher-than-expected cumulative frequencies (Figure 2A). We calculated pairwise repertoire overlap distance between individuals based on the Jaccard index (see Methods). Using this distance, we performed hierarchical clustering (Figure S1A,B) and found that individuals clustered by age and repertoire diversity (Figure 2A), especially when looking at superpublic CDR3 sharing patterns. Indeed, upon splitting the dendrogram of superpublic CDR3s into four clusters (Figure 2B), we found that individuals in the different clusters had different repertoire sizes and age distribution (Figure 2C,D). Clusters #2 and #4 showed maximum divergence: individuals in cluster #2 had an average of 0.6 x10^6^ different CDR3 sequences and a mean age of 40, against only 0.1 x10^6^ sequences and a mean age of 93 in cluster #4 (Figure 2C,D). We wondered whether variations in TDT activity with age (Bonati et al., 1992; Deibel et al., 1983; Pahwa et al., 1981) could impact on repertoire sharing. When we aligned each CDR3 to the germline from the reference genome and counted the number of mismatches (see Methods), we found that indeed, cord blood CDR3s (TDT-negative) contained fewer mismatches than samples from other age groups (Figure 2E). Finally, when we grouped CDR3aa by descending order of frequency (see Methods), we found that the most frequent CDR3aa displayed fewer mismatches than those with lower frequency, most distinctively in cord blood (Figure 2F).

**Figure 2:**
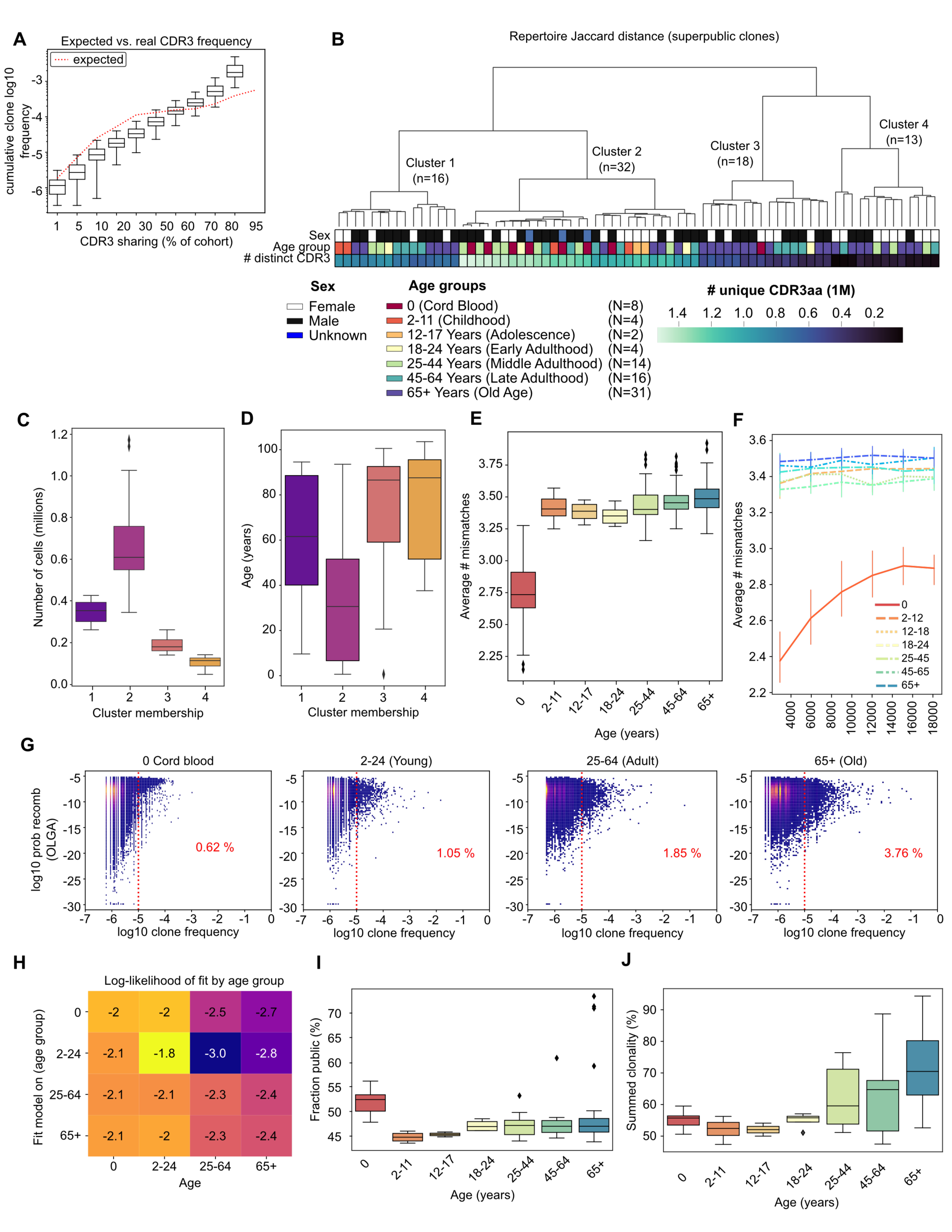
CDR3 sharing between individuals as a function of age. **(A)** Cumulative log10 frequency of CDR3 binned by cohort sharing percentage. The red dotted line indicates the expected frequency given the number of individuals in each sharing bin. **(B)** Hierarchical clustering of individual repertoires based on pairwise Jaccard distance. Dendrogram leaves representing individuals are colored by sex, age group, and the number of distinct CDR3aa found in individual repertoires. **(C and D)** Boxplots show the distribution of the number of distinct public CDR3aa found in individuals and the age of individuals in each of the four clusters from Figure 2B. **(E)** The median number of mismatches to the germline found in CDR3aa of individuals by age group. **(F)** The average number of mismatches to the germline found in CDR3aa of individuals grouped by CDR3aa frequency in each repertoire. Colors represent various age groups. **(G)** Aging correlates with an accumulation of high-frequency clonotypes with a high recombination frequency (as determined by OLGA score). **(H)** Mean log-likelihood of fit for CDR3aa found in age groups in abscissa for models trained on age groups in ordinate. Each row represents a model trained on the age group, and each column represents the test CDR3aa; each cell contains the mean log-likelihood of fit for a Gaussian mixture model (see Methods). **(I)** The proportion of public CDR3aa in different age groups. **(J)** Summed clonality of public CDR3aa in various age groups.

Further clone size analyses showed that as individuals age, they accumulate in their repertoires more very high-frequency CDR3aa (at frequencies above 0.0001 of total repertoire), which have a recombination frequency in the higher ranges (above 10^-10^) (Figure 2G). Moreover, a low clonal frequency for high recombination frequency sequences could be partly due to undersampling in the repertoire (Sethna et al., 2019) since only a certain amount of CDR3s are sequences. We used a two-step strategy to evaluate the relationship between clonality and the probability of recombination at different ages. We fitted a Gaussian mixture model for each age group, and then we calculated the log-likelihood of data from other age groups under this model (see Methods). We found that models fitted on repertoires of younger individuals did not fit with data from older individuals. However, since models fitted on older individuals had a similar likelihood for all age groups, we concluded that older repertoires retain characteristics of younger repertoires and outgrow them with time (Figure 2H). What distinguishes older repertoires from younger ones is a large quantity of high-frequency (presumably hyperexpanded) CDR3aa with a high recombination probability (Figure 2G,H).

For individual samples in the Britanova cohort, the proportion of public CDR3aa was maximum in cord blood, dropped abruptly in children, and increased progressively with age after that (Figure 2I). As a result, the proportion of public CDR3aa in subjects ≥ 65 years of age was similar to that in cord blood. The progressive increase in the fraction of public CDR3aa from childhood to old age was even more conspicuous when considering the clonality of each CDR3aa (see Methods, the section on CDR3 sharing): almost 70% of repertoires in individuals ≥ 65 years of age were composed of public CDR3aa (Figure 2J). Though cord blood and samples from subjects ≥ 65 years of age contained similar proportions of public CDR3aa (Figure 2I), their clonality was very different (Figure 2J). Cord blood cells had a more uniformly distributed repertoire of public CDR3aa, without the hyperexpanded clones present in subjects ≥ 65 (Figure 2I-J). We validated our observation in two additional cohorts. In the Emerson cohort, containing TCR-Seq data from 666 healthy individuals (Emerson et al., 2017), we could split individuals by CMV status. We found that an age-related skew in public fraction can be observed in CMV+ and CMV- subjects (Figure S2A-D). The Thome cohort is smaller but contains TCR-Seq data from deceased donors’ spleen and lymph nodes rather than blood (Thome et al., 2016). T cells were sorted by naive or effector memory phenotype in this study; we, therefore, analyzed those categories separately. We found the same trend of sharing by age group for the naive T cells (Figure S2E-F) but not for the effector memory T (TEM) cells in secondary lymphoid organs (Figure S2G-H). The latter divergence warrants further investigation but must be considered preliminary because it is based on analyses of a small cohort of deceased donors.

These results indicate that as individuals age, their repertoire becomes preferentially populated by clones with high recombination frequencies. A high recombination frequency is likely instrumental in the abundance of highly public clones. Another possible explanation could be a preferential expansion of these T cells due to homeostatic proliferation (Murray et al., 2003) or immune activation. To test the latter hypothesis, we used the ERGO software (Springer et al., 2020) to predict the recognition of a large set of HLA-associated peptides recognized by public and private CDR3aa. Our dataset included 25,270 human peptides (Pearson et al., 2016) and 20,961 viral-derived peptides (Vita et al., 2019). We found that peptides recognized by shorter and more shared CDR3aa had higher TCR binding scores than peptides recognized by longer and less shared CDR3aa (Figure S3A-C). For further validation, we used single-cell TCR sequencing analyses of T cells responding to 50 different HLA-associated peptides (see Methods). We labeled CDR3aa that recognized more than one peptide as polyreactive. The highly polyreactive CDR3aa had an average length ranging from 10 to 17 amino acids (Figure S3D), had fewer mismatches (Figure S3E), and a higher recombination probability (Figure S3F), consistent with results from (Lu et al., 2019). One caveat of these data is the limited number of peptides tested. Nevertheless, they highlight polyreactivity as another characteristic of shared CDR3aa and suggest that polyreactivity contributes to the high clonality of public CDR3aa in older individuals.

### The impact of sex on the TCR repertoire

Aside from age, male sex is the factor with the most negative impact on thymic output (Clave et al., 2018). Therefore, we analyzed the potential influence of sex on CDR3 repertoire diversity and publicness by grouping individuals into broader age groups to maintain adequate comparison numbers between categories (Figure S3J). Overall, we found that males had fewer CDR3aa in their repertoires than females: this was the case for public (Figure 3A) and superpublic CDR3aa (Figure 3B). We then analyzed a set of almost 160,000 high-confidence SARS-CoV2-specific TCRs (Nolan et al., 2020). Since the Britanova cohort was sequenced in 2014, all individuals were unexposed to SARS-CoV2. Notably, we found a high number of SARS-CoV2-specific CDR3aa in the Britanova cohort repertoires (Figure 3C). Again, males had fewer SARS-CoV2- specific CDR3aa in their repertoires than females (Figure 3C). When we examined repertoire diversity using Shannon entropy, we found that repertoires of females were more diverse than those of males (Figure 3D-F). Accordingly, small-size CDR3aa clonotypes represented 70% of the repertoire in females and 50% in males (Figure 3 G,H). In contrast, hyperexpanded CDR3 clonotypes constituted 30% of repertoire in males and 10% in females. Differences between males and females were present in all age groups and always reached statistical significance in subjects aged 2-45 but not in other groups (Figure 3A-F). These results highlight a prominent sexual dimorphism in the TCR repertoire and suggest that it results from differences in thymic output. Female repertoires are more diverse, and males present a lower measured repertoire diversity with hyperexpansion of selected TCR clonotypes.

**Figure 3:**
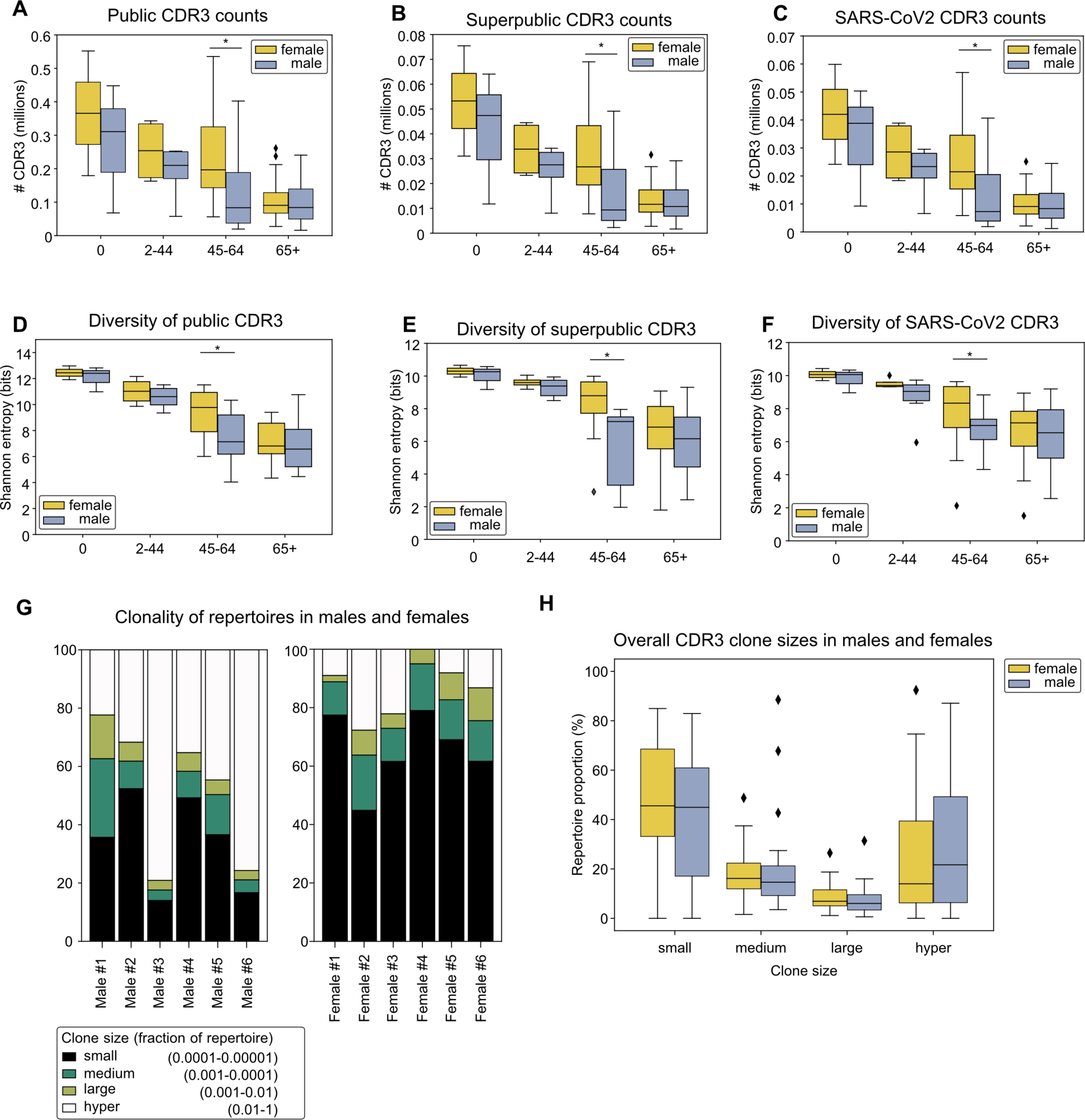
CDR3 sharing between individuals as a function of sex. Absolute numbers of **(A)** public, **(B)** superpublic, and **(C)** SARS-CoV2-specific CDR3aa in male and female individuals by broad age groups. Difference statistically significant (p<0.05) for young individuals (ages 2-44). Shannon entropy for **(D)** public, **(E)** superpublic, and **(F)** SARS- CoV2-specific CDR3aa in males and females. Statistically significant differences (p<0.05, Mann-Whitney-Wilcoxon) for subjects aged 2-44 and 45-65. **(G)** Clonality of CDR3aa in repertoires of adult males and females, binned by clone size. **(H)** Boxplot showing the distribution of overall clone sizes of CDR3aa in males and females of all age groups.

### Sharing of disease-specific CDR3s in different age groups

Prompted by the results of our analyses of SARS-CoV2-specific CDR3s, we downloaded and explored the McPAS database, a manually curated catalog of pathology-associated TCR sequences (Tickotsky et al., 2017). We found minimal overlap (0.1-3%) between TCRs in two McPAS categories: microbial pathogens and autoimmune diseases (Figure S4A). To gain further insight into disease-related CDR3s, we took CDR3aa listed in the McPAS microbial pathogens dataset and analyzed their frequency in subjects from the Britanova cohort (Figure 4A). The hierarchical clustering dendrogram was separated into three clusters for individuals (I1 to I3) and five clusters for CDR3aa (C1 to C5). Age had a dramatic influence on both dimensions of this orthogonal clustering. Among clusters for individuals, cluster I2 was composed solely of cord blood samples, whereas individuals in clusters I1 and I3 had a mean age of 82 and 26 years of age, respectively (Figure 4B). The CDR3aa-based clustering adopted the following pattern: i) CDR3aa in cluster C1 were present almost exclusively in cord blood, ii) those in cluster C2 were present in few individuals without any clear pattern, and iii) CDR3aa in clusters C4 and C5 were present in young individuals (cord blood and <45 y.o.) (Figure 4A). Cluster C3 was remarkable in that it contained the most highly shared CDR3aa; they were found at high frequency in cord blood and lower frequency in almost all other individuals. CDR3aa in cluster C3 were shorter and displayed a greater recombination frequency than CDR3aa in the four other clusters (Figure 4C,D). Observations on microbial pathogens-related CDR3aas were replicated in autoimmune disease-associated CDR3aa (Figure S4B-E). First, cord blood (cluster I1 in Figure S4B) contained more autoimmunity-associated CDR3aa. Second, the most highly shared CDR3aa (cluster C1 in Figure S4B) were shorter and displayed a greater recombination frequency than CDR3aa in the four other clusters.

**Figure 4:**
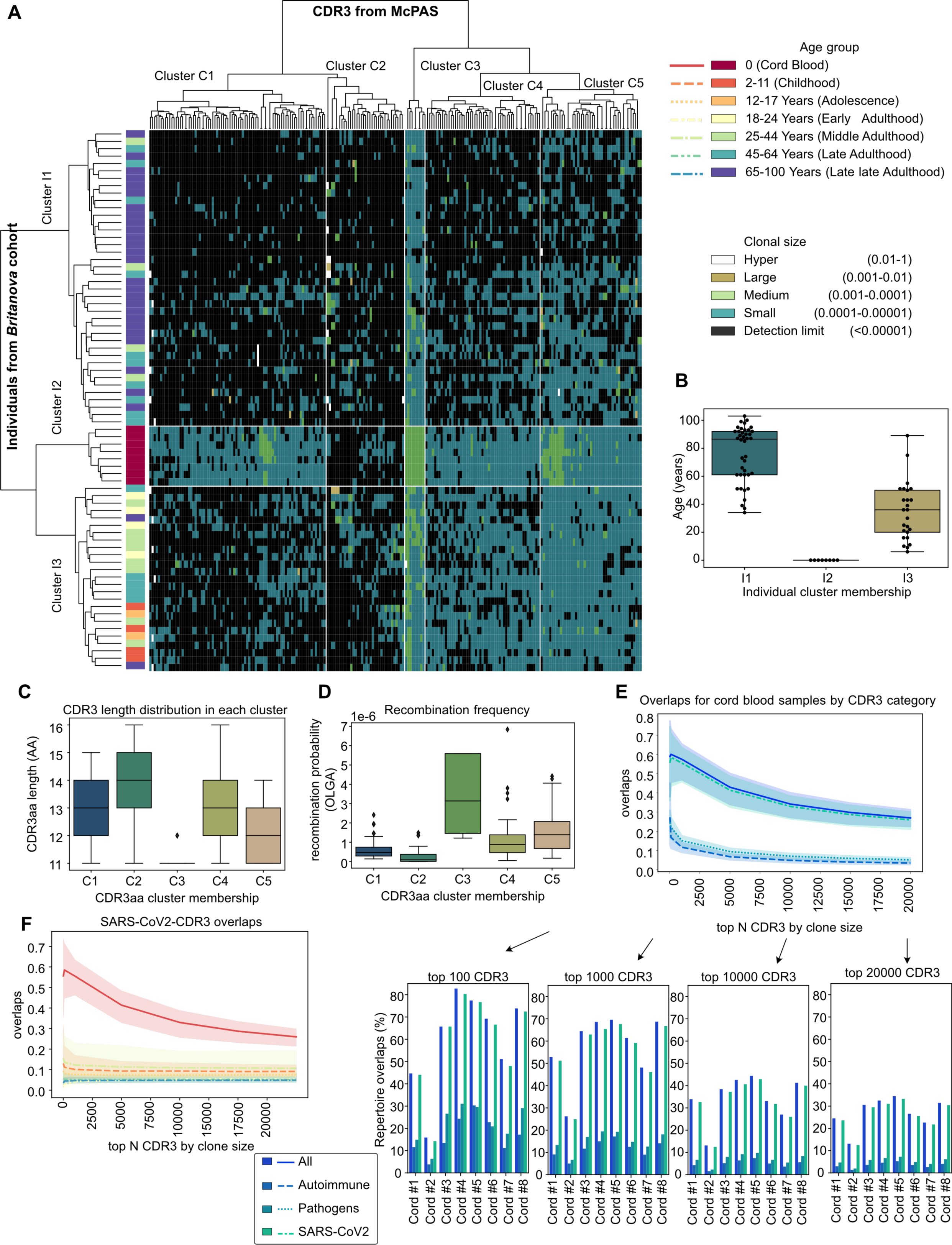
Cord blood samples contain pathology annotated CDR3s. **(A)** Heatmap shows, for subjects of the Britanova cohort, the frequency of CDR3aa listed in the McPAS microbial pathogens dataset (Tickotsky et al., 2017). Rows represent individuals, columns unique CDR3aa, and cell color indicates CDR3aa clone size. Row dendrogram leaves are colored by age group. **(B)** Age distribution for individuals in three individual (Y-axis) clusters from (A). **(C)** Boxplot showing CDR3 lengths for CDR3 in the five X-axis clusters from (A). **(D)** Boxplot showing predicted recombination frequency for CDR3 in five X-axis clusters from (B). **(E)** Line and barplots show the percentage CDR3aa responsive to autoantigens (Tickotsky et al., 2017), SARS-CoV2 (Nolan et al., 2020), or other pathogens (Tickotsky et al., 2017) in individual cord blood samples from the Britanova cohort, by varying top N most frequent CDR3aa. **(F)** Line plots showing the percentage of SARS-CoV2-specific CDR3aa among the top N most frequent CDR3aa. Line colors and types correspond to age groups as in panel A.

The key finding was that almost all disease-related CDR3aas were found in cord blood. Indeed, 18 to 75% of CDR3aa in individual cord blood samples (Britanova cohort) were responsive to SARS-CoV2 [i.e., present in high-confidence SARS-CoV2-specific TCRs (Nolan et al., 2020)] (Figure 4E). A significant proportion of the most frequent clones in cord blood CDR3aa was also responsive to other pathogens and autoantigens (Figure 4E). The large size of the SARS-CoV2-specific TCR dataset explains why more cord blood CDR3aa appeared responsive to SARS-CoV2 than other pathogens and autoantigens. Moreover, since the Britanova cohort only contains CDR3 beta sequences, without CDR3 alpha or HLA, this result is likely an overestimation but can still be used to compare between age groups and individuals. In some cord blood samples, the summed proportions of CDR3aa responsive to autoantigens, SARS-CoV2, and other pathogens were superior to 100% (Figure 4E). This is most likely justified by the polyreactivity of public TCRs (Figure S3). We conclude that all individuals have many disease-reacting clones at a high frequency in their repertoires before birth. Are disease-related CDR3s present in older subjects? To address this question, we calculated the percentage of disease-related CDR3aa present in top N CDR3aa from individuals of various age groups (Figure 4F, S5A-C). Two points can be made from this analysis. First, in cord blood, disease-related CDR3aa are enriched in high-frequency clonotypes. Second, the remarkable representation of disease-related CDR3aa in the “pre-immune” repertoire of cord blood is lost in older individuals.

Our data support the notion that TCRs generated during fetal life can persist (or be continuously generated) for decades in adults (Pogorelyy et al., 2017). More importantly, they show that most of these TCRs participate in a wide variety of immune responses in adult life. Globally, our data presented so far suggest the existence of two types of CDR3: the superpublic ones, shared by many individuals and present before birth, and the private repertoire, dependent on TDT modifications. For the remainder of the study, we will refer to these two types of TCRs as *neonatal* and *TDT-dependent*.

### Negative selection targets TDT-dependent TCRs

Irrespective of their TCR type, neonatal or TDT-dependent, T cells are subjected to intrathymic positive and negative selection. Mutations in the AIRE protein are known to be associated with perturbations in negative selection and thereby causing autoimmunity (Liston et al., 2003). We, therefore, analyzed the CDR3s of the Sng cohort, which contains subjects with AIRE mutations and healthy controls (Sng et al., 2019). We found that CDR3aa repertoires of AIRE-mutated individuals had a lower public fraction than healthy repertoires (Figure 5A), with a lower recombination frequency (Figure 5B), a higher number of mismatches per CDR3aa (Figure 5C) for both regulatory and conventional T cell compartments, and longer CDR3aas (Figure 5D). These results point towards enrichment in TDT-dependent CDR3s in repertoires of AIRE- mutated individuals, which in turn suggests that thymocytes with TDT-dependent TCRs are prime subjects of negative selection.

**Figure 5:**
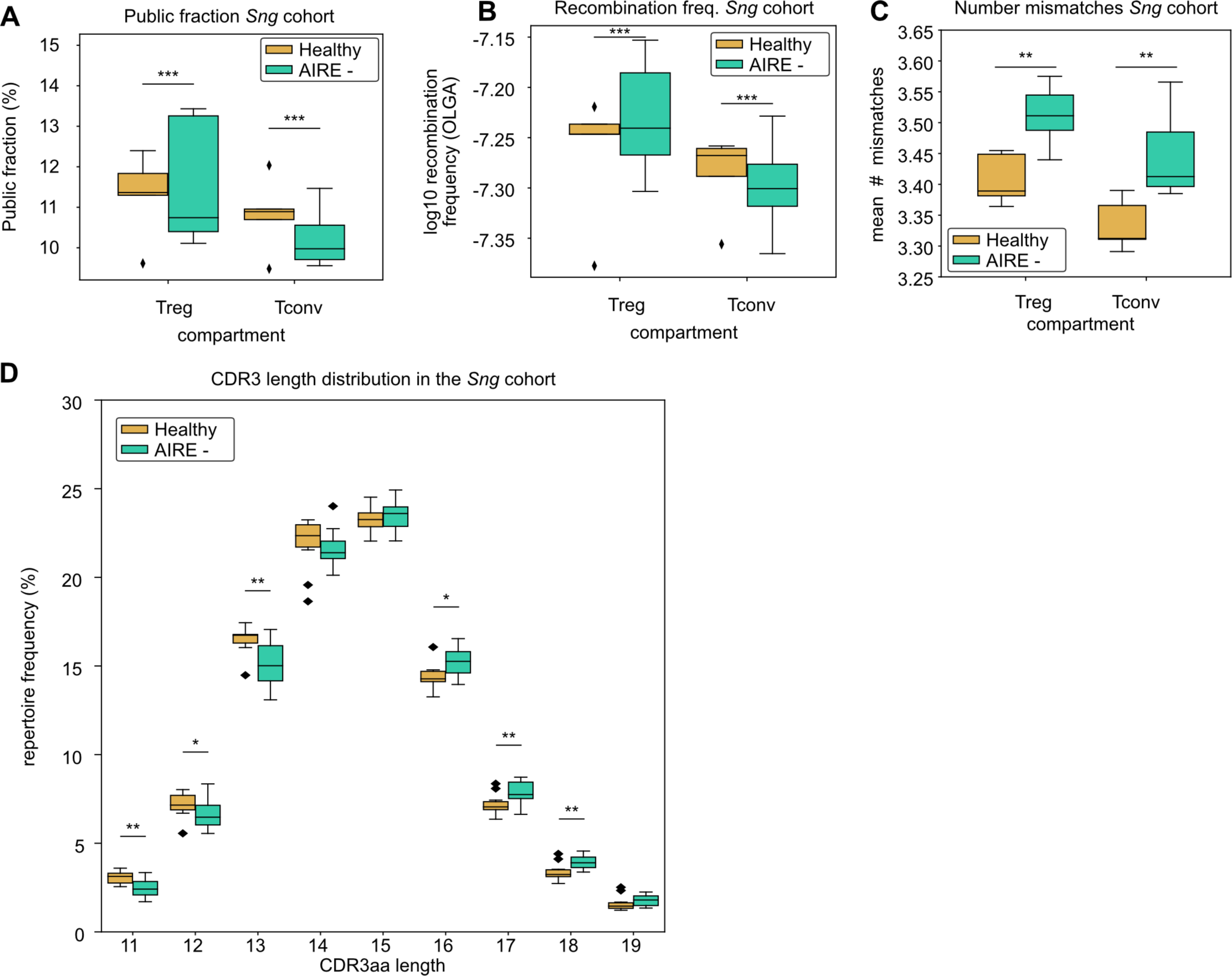
CDR3aa profile in subjects with AIRE mutations. Boxplots showing, for the regulatory (Treg) and conventional T cell (Tconv) compartments, **(A)** the public fraction, **(B)** the recombination frequency, and **(C)** the number of mismatches in subjects with AIRE mutations vs. controls. **(D)** Boxplots show the CDR3aa length distribution in the Sng cohort. (*p<0.05, ** p< 0.01, ***p<0.001, Mann-Whitney-Wilcoxon).

### Effect of the TCR repertoire on graft-versus-host disease

To further evaluate the potential impact of the two types of TCRs, we reasoned that the best strategy would be to use a model in which the readout depends exclusively on T cells. Acute graft-versus-host disease (aGVHD) following allogeneic hematopoietic cell transplantation (AHCT) represents such a model. Indeed, donor T cells, particularly the CD4+ subset, are necessary and sufficient for the occurrence of aGVHD (Ni et al., 2017; Socié and Blazar, 2009). They initiate aGVHD via recognition of host alloantigens (Martin et al., 2017; Vincent et al., 2011). Therefore, we analyzed TCRs in purified CD4+ T cells from 73 AHCT donors. Donors and recipients were HLA-matched siblings. The T cells were obtained from the peripheral blood of donors on the day of transplantation and submitted to RNA sequencing. To extract CDR3 sequences from RNA sequencing reads, we used the MIXCR software (Bolotin et al., 2015; Li et al., 2017). We classified donors as aGVHD+ or aGVHD-, depending on whether their recipient presented or not severe aGVHD (see Methods). Notably, aGVHD+ donors had lower CDR3 diversity than aGVHD- grafts (Figure 6A). We used a treemap to display both diversity and clone size in two representative donors. Treemaps offer a visual representation of diversity at a glance, and we used these plots to compare two representative examples of aGVHD- and aGVHD+ donor repertoires. In the aGVHD+ donor, three hyperexpanded clones occupied almost ⅓ of the repertoire (Figure 6B), while the aGVHD- donor did not have this skew (Figure 6C). The CDR3aa in aGVHD+ grafts were longer (Figure 6D), had a lower recombination frequency, and more numerous mismatches than CDR3aa in aGVHD donors (Figure S5D-E). We then split the cohort by the median or quartiles and generated Kaplan-Meier curves to assess the impact of CDR3 features on the occurrence of aGVHD (Figure 6E-H, Figure S6). Overall, grafts containing a higher proportion of CDR3 with neonatal features caused less aGVHD. These features were: CDR3 length in amino acids (Figure 6E), percentage overlap with cord blood samples (Figure 6F), recombination frequency (Figure 6G), and Simpson diversity index (Figure 6H).

**Figure 6:**
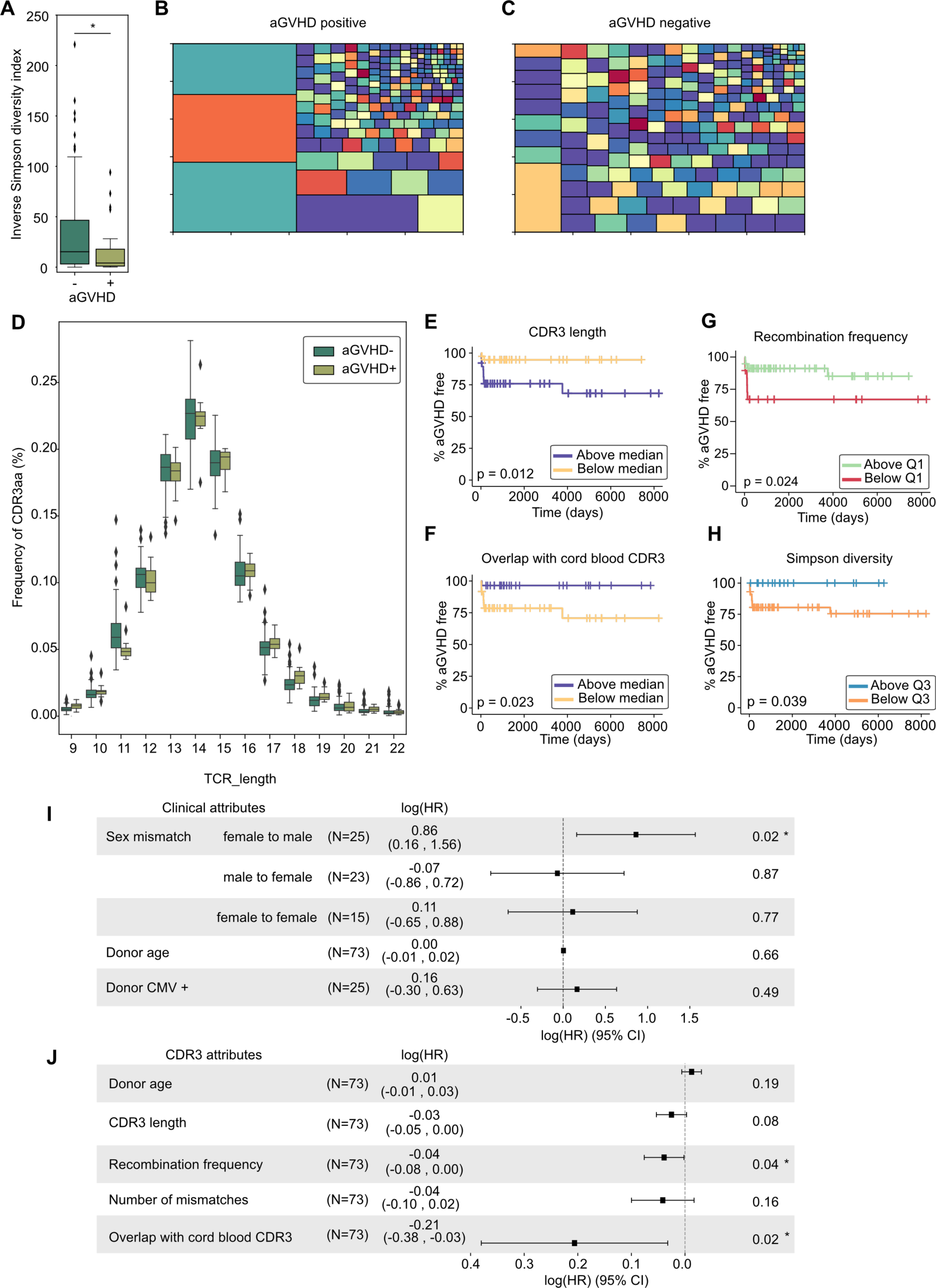
CDR3aa in CD4 T cells from aGVHD+ and aGVHD- AHCT donors. **(A)** Inverse Simpson diversity index of CDR3 repertoires found in aGVHD+ and aGVHD- grafts. Treemaps showing CDR3aa diversity and clone sizes for two representative donors **(B)** aGVHD+ and **(C)** aGVHD-, colors selected at random for better visual distinction. **(D)** CDR3aa length distribution in aGVHD+ and aGVHD- donors. Kaplan-Meier curves representing aGVHD onset for grafts split into two groups by median or quantiles according to **(E)** CDR3 length, **(F)** overlap with cord blood CDR3aa, **(G)** recombination frequency, and **(H)** Simpson diversity index. CoxPH models calculating hazard ratios for **(I)** clinical characteristics and **(J)** CDR3 repertoire of the donor. (*p<0.05, **p<0.01, ***p<0.001).

Finally, we used Cox proportional hazards (CoxPH) models to evaluate more accurately the impact of clinical and CDR3 features on the risk of aGVHD. For the clinical characteristics model, the sole significant correlation was a higher rate of aGVHD in male recipients of female grafts (Figure 6I). These results are concordant with previous reports (Kim et al., 2016). For the CDR3 model, we found that a high number of neonatal CDR3 and a high average recombination frequency decreased the risk of aGVHD (Figure 6J). Other characteristics and clinical traits such as donor age and CMV status had no significant impact (Figure 6I,J). Collectively, these results strongly suggest that donors with a higher proportion of neonatal TCRs cause less aGVHD and that aGVHD is initiated primarily by TDT-dependent TCRs.

### A stratified model of the TCR repertoire

Our final goal was to evaluate the importance of discrete features in defining neonatal and TDT-dependent TCRs. Our reasoning was based on two assumptions. First, we assumed that cord blood samples contained exclusively neonatal TCRs while all other age groups contained a mix of neonatal and TDT-dependent TCRs. Second, since thymic output and TDT activity reach their zenith during childhood, we postulated that children would generate the greatest diversity of TDT-dependent TCRs. Therefore, to get a pure and diversified population of TDT-dependent CDR3s, we selected CDR3s present in children but not in cord blood. We then confirmed that, compared to neonatal CDR3s, the TDT-dependent CDR3s were longer (Figure 7A) had more mismatches and a lower recombination probability (Figure 7B-C). Notably, they also displayed a different V and J gene usage (Figure 7D-E).

**Figure 7:**
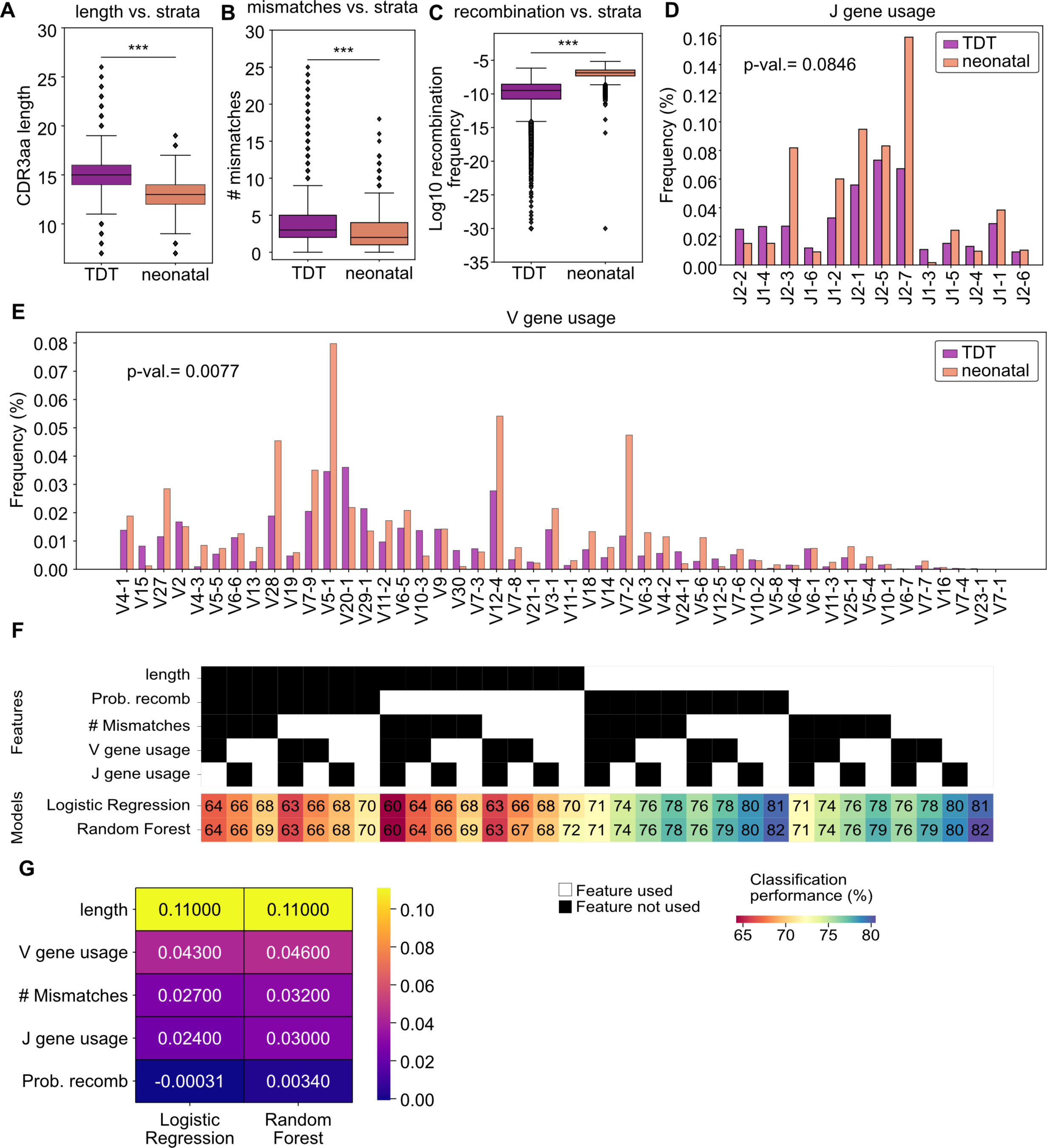
Features of neonatal vs. TDT-dependent TCRs. Boxplots depict **(A)** the CDR3aa length, **(B)** the median number of mismatches to the germline, and **(C)** the median log10 recombination frequency of the TDT-dependent and neonatal strata. **(D)** J gene and **(E)** V gene usage frequencies for CDR3aa in TDT-dependent and neonatal strata. **(F)** Feature ablation study showing classification accuracy on held-out data for each feature combination. Black/ white squares signal exclusion/inclusion of features in the dataset, and the color scheme shows classification performance. **(G)** Coefficients of the linear model fitted on feature ablation study (see Methods). (*p<0.05, **p<0.01, ***p<0.001, Mann-Whitney- Wilcoxon).

On this dataset, we trained a logistic regression model and random forest to verify if the nonlinearity of the model could have an impact on the performance. Using all the five features (recombination frequency, # mismatches, CDR3 length, V gene, J gene), we performed an ablation study by obtaining all possible combinations of presence/absence, totaling 31 combinations of features (Figure 7F). We trained the two models on the dataset for each combination and evaluated their performance on a held-out CDR3 repertoire of each type (the entire individual’s repertoire). The performance of each model on the held-out data is represented as a single column, where black squares symbolize the absence and white squares the presence of a feature, and the performance squares are colored by the percentage of accuracy of classification (Figure 7F). The CDR3 length was crucial to the model; without the CDR3 length, the model’s performance was close to the baseline of 60%, which is the proportion of neonatal CDR3s in the dataset. Adding the length improves classification accuracy by about 10% for all conditions. Numbers of mismatches and V/J gene usage had a more modest effect on the performance, with an accuracy gain of about 5% each. Moreover, V and J gene usage was non-redundant and having both yielded better performance than only having one or the other for both models. The inclusion of the recombination frequency did not impact the performance, most likely because it is largely redundant with CDR3 length (Figure S7A).

Finally, to validate the order of importance of the features, we fit a linear regression model on the presence/absence of features (see Methods). This allowed for the direct comparison of the relative importance of the features based on the coefficients assigned to each feature (Figure 7G). We found the following importance hierarchy: length > (# mismatches, V/J gene usage) > probability of recombination (Figure 7G). Consistency between the logistic regression and the random forest models suggests that these features robustly discriminate between neonatal and TDT-dependent TCRs. We then used the trained model to classify each CDR3 in the cohort and found that, as expected, there is a considerable dip in the proportion of neonatal TCRs after birth (Figure S7B). Then, from infancy to adulthood, there is a progressive increase in the proportion of neonatal clonotypes (Figure S7B), probably because of their polyreactivity (Figure S3). Afterward, the proportion of neonatal CDR3 remains relatively stable with a slight trend downwards with advancing age (Figure S7B).

## DISCUSSION

This report analyzed the amino acid sequence of over 100 million TCR CDR3 beta chains from over 1,000 subjects. Since both TCR chains contribute to antigen specificity, we would have wished to extend our analyses to CDR3aa alpha/beta pairs obtained through single-cell studies (Pauken et al., 2022). However, there is no available dataset that contains paired CDR3 alpha and beta chains from large human cohorts. Indeed, because of methodological constraints inherent to single-cell approaches, paired CDR3 alpha and beta chain sequencing has been limited to small numbers of subjects, typically 1 to 15 (Fischer et al., 2020; Pauken et al., 2022; Tanno et al., 2020). Focusing on CDR3 amino acid sequences instead of nucleotides and V/J gene usage allowed us to uncover more numerous public and superpublic CDR3aa than anticipated, since this analysis considers synonymous codons. In a way, our analysis examines the functional features of TCRs, which are determined by their amino acid sequence. We found stark differences between male and female repertoires, as well as age-specific and disease-specific repertoire features.

Age and sex are associated with important differences in immune responses to pathogens and self-antigens (Brodin and Davis, 2017; Liston et al., 2016). Thymic involution is instrumental in decreasing immunocompetence with age and represents a major public health issue, as illustrated by the COVID pandemic (Mittelbrunn and Kroemer, 2021; Palmer et al., 2018; Yousefzadeh et al., 2021). Aside from age, male sex is the factor with the most negative impact on thymic output (Palmer et al., 2018). We report that both aging and male sex are associated with decreased TCR diversity and hyperexpansion of public clonotypes. Female TCR repertoires are more diverse, and males compensate for their lower repertoire diversity via hyperexpansion of selected TCR clonotypes. These data argue for a strong mechanistic link between thymic output and TCR diversity.

Analyses of cord blood samples were particularly instructive. In the absence of TDT, TCRs produced before birth have short CDR3s, few mismatches (relative to germline sequences), and a biased V/J gene usage. These neonatal TCRs persist (or are continuously replenished) throughout life, are highly shared among subjects, and are polyreactive to self and microbial HLA-associated peptides. Three factors likely contribute to the large clone size and extensive sharing of neonatal TCRs over a lifetime. First, they have a high recombination frequency; in other words, they are easy to assemble during V(D)J recombination. Second, their high reactivity to self-antigens (Figure S2A) should theoretically favor their positive selection in the thymus and their homeostatic proliferation in the periphery (Ernst et al., 1999; Hogquist and Jameson, 2014). Third, our analyses of subjects with AIRE mutations revealed that neonatal TCRs were less affected by negative selection in the thymus than TDT-dependent TCRs. Thus, neonatal TCRs may integrate all the “Goldilocks” conditions for intrathymic selection and survival in the periphery. Notably, polyreactivity to self-antigens could also favor the commitment of thymocytes bearing neonatal TCRs toward either the regulatory or alternative T cell lineages (Sood et al., 2021; Vrisekoop et al., 2014). This possibility should be explored in future studies.

While both TCR chains as well as the MHC molecule contribute to antigen specificity, in practice, most analyses of the T-cell repertoire have focused on CDR3 beta, mainly for two reasons. Firstly, the sequencing of CDR3 beta is more robust than that of CDR3 alpha (Barennes et al., 2020). Secondly, CDR3 beta is the main contributor to TCR antigen specificity (Springer et al., 2020, 2021). Accordingly, though the prediction of antigen specificity is improved by paired CDR3 alpha/beta sequencing, predictions based only on CDR3 beta perform well (Fischer et al., 2020). Furthermore, analyses of CDR3 alpha and beta chain pairing in close to 1 million clonotypes from 15 individuals (Tanno et al., 2020) support our main conclusions: shared CDR3aa are relatively short with few TDT-dependent additions (Tanno et al., 2020).

The effect of the HLA genotype on the repertoire of CDR3 beta sequences is detectable (Khosravi-Maharlooei et al., 2019; Tanno et al., 2020) but remains reatively small (Emerson et al., 2017; Heikkilä et al., 2021; Pogorelyy et al., 2018; Springer et al., 2021). This is explained at least in part by the fact that most MHC-associated peptides can bind to multiple HLA alleles. Consistent with our results, studies in mice revealed that the most highly shared TCRs among mice with different MHC genotypes have shorter CDR3 sequences (Lu et al., 2019). In contrast, a single TCR has been shown to be capable to bind with as many as million different antigens (Bentzen et al., 2018; Natarajan and Krogsgaard, 2018; Wooldridge et al., 2012; Zhang et al., 2018). Therefore, while our analysis only includes beta chains and therefore overestimates polyreactivity, we found it remarkable that 10-80% of CDR3s in cord blood samples were reactive to SARS-CoV2, other pathogens, or autoantigens (Figure 4E. (Figure 4E). This means that humans are born with a TCR repertoire that can have a lifelong influence on their response to pathogens and the risk of autoimmunity. From an evolutionary perspective, the size of human populations has been limited by the rate of infant mortality. Hence, it would seem convenient to be born with a polyreactive T cell repertoire responsive to common pathogens.

In contrast to neonatal TCRs, TDT-dependent TCRs are longer, less shared, contain more mismatches, and display a different V/J gene profile. Their production is maximal during infancy, when thymic output and TDT activity reach a summit, and slowly decreases after that. We found that TDT-dependent TCRs were more abundant in subjects with AIRE mutations. This suggests that negative selection preferentially eliminates TDT-dependent TCRs. The ultimate role of TDT remains unclear. By ultimate role, we mean the evolutionary selected biological advantage conferred by TDT. In mice, deletion of TDT does not increase susceptibility to pathogens or the incidence of autoimmunity but decreases the breadth of anti-viral responses (Haeryfar et al., 2008; Kedzierska et al., 2008). However, for the immune system, evolutionary convergence towards a higher diversity is thought to be a protection mechanism to get ahead of the arms race with pathogens (Liston et al., 2021). Therefore, a plausible hypothesis is that the presence of TDT-dependent TCRs confers an additional, more “private” layer of security against the emergence of antigen-loss variants.

aGVHD is a harbinger of chronic GVHD and has remained the nemesis of patients and physicians during the entire history of AHCT, partly because its occurrence is unpredictable. Our aGVHD cohort was composed of HLA-matched siblings. In this situation, aGVHD is caused by donor T cells that react against host minor histocompatibility antigens (Vincent et al., 2011; Warren et al., 2012). On the other hand, histoincompatibility does not always elicit fatal GVHD. Indeed, in patients that received AHCT from donors presenting multiple disparities for minor histocompatibility antigens, only 73% developed aGVHD (Martin, 1991). It has been hypothesized that some AHCT donors might be stronger alloresponders than others (Baron et al., 2007). In our cohort of 73 donor-recipient pairs, the occurrence of severe aGVHD was strongly associated with a low proportion of neonatal TCRs in the donor repertoire. Such a protective effect of neonatal TCRs would explain reports that AHCT with cord blood rather than adult hematopoietic cells may be associated with a lower risk of GVHD (Cohen et al., 2020). Moreover, while some studies report no relationship between post-transplant diversity and GVHD occurrence (Buhler et al., 2020), our results on the diversity in grafts (pre-transplant) are consistent with those of Yew and colleagues, who report that a lower TCR diversity was correlated with GVHD occurrence and relapse, while a higher percentage of cord-blood cells was correlated with a higher repertoire diversity (Yew et al., 2015). If our observation is validated in further studies, it will justify the preferential selection of AHCT donors with a high proportion of neonatal TCRs in their peripheral blood.

Together, our data support an emerging model in which the T cell repertoire is composed of two strata with differential reactivity to self and non-self antigens: public neonatal TCRs and private TDT-dependent TCRs. This model is remarkably coherent with insightful theoretical predictions by Vrisekoop and colleagues who labeled the two strata the “somatic” repertoire and the “ur”-repertoire (Vrisekoop et al., 2014). Our model is also consistent with functional studies demonstrating that neonatal T cells can no longer be considered immature versions of adult cells. On the contrary, they are highly functional and respond rapidly to antigenic challenges (Davenport et al., 2020; Rudd, 2020).

## Author Contributions

A.T., C.P. and S.Le. designed the study. A.T. performed the main <u>bioinformatic</u> analyses and result interpretation. J.S., J.-P.L., S.La., L.B., S.Le. and A.B. performed <u>RNA Sequencing</u> experiments. P.B., J.-D.L. and G.E. contributed to bioinformatic analyses. A.T., P.B., J.-D.L., G.E., S.Le. and C.P. contributed to the analysis and interpretation of data and results. A.T. and C.P. wrote the manuscript and all authors edited and approved the final manuscript.

## Acknowledgments

This study was supported by grant FDN-148400 from the Canadian Institutes of Health Research (to C.P.). A.T. was supported by a studentship from the Canadian Institutes of Health Research, and J.D.L. by a studentship from the Fonds de Recherche Québec – Santé.

## METHODS

### Resource Availability

#### Lead contact

Further information and requests for resources should be directed to and will be fulfilled by the lead contact, Claude Perreault.

### Data and code availability

RNA-Seq data from the GVHD cohort has been deposited in the Sequence Read Archive and are publicly available at BioSample accession PRJNA832136. DOIs are listed in the key resources table.

All original codes generated during this study are available in the form of python jupyter notebooks on Github, at the address listed in the key resources table.

Any additional information required to reanalyze the data reported in this paper is available from the lead contact upon request.

### Experimental Model and Subject Details

#### GVHD cohort individuals

The *GVHD* cohort included 73 healthy sibling-matched donors. Written informed consent was obtained from all patients or their legal guardians before sample collection or hematopoietic stem cell transplantation. For each sample, peripheral blood mononuclear cells (PBMC) were collected, and CD4 T cells were isolated by immunomagnetic positive selection (EasySep Human CD4 positive selection kit; Stemcell Technologies). Sex and age for these individuals can be found in the metadata corresponding to the RNA-Seq data available from the Sequence Read Archive under accession number PRJNA832136.

### Method Details

#### TCR sequencing datasets

We downloaded TCR sequences and additional data from four non-overlapping cohorts: (Britanova et al., 2016; Emerson et al., 2017; Sng et al., 2019; Thome et al., 2016). A total of 980 human subjects were included in these cohorts, with 401 females and 517 males; the sex of 47 subjects was unknown. For further details on numbers of sequences, see Supplementary Figure S3H.

#### Combined TCR-Seq/single-cell RNA-Seq data from antigen-specific CD8+ T cells

We obtained from the 10x Genomics data repository V(D)J sequence information generated by 10x Genomics CellRanger for 150,000 CD8+ T cells isolated from four healthy donors. Data was downloaded from (https://www.10xgenomics.com/welcome?closeUrl=%2Fresources%2Fdatasets&lastTouchOfferName=CD8%2B%20T%20cells%20of%20Healthy%20Donor%201&lastTouchOfferType=Dataset&redirectUrl=%2Fresources%2Fdatasets%2Fcd-8-plus-t-cells-of-healthy-donor-1-1-standard-3-0-2). We did the following data pre-processing. We filtered out barcodes associated with non-unique cells or did not have a resolved CDR3 amino acid or nucleotide sequence. As previously reported (Schattgen et al., 2021), we found substantial non-specific binding in donor 1 and excluded this sample.

#### GVHD cohort individuals

RNA was extracted from purified CD4 T cells by Trizol-Column (PureLink RNA Mini kit; Thermo Fisher Scientific). RNA was quantified by U.V. spectrophotometry (Tecan Infinite M1000), and quality was verified by Bioanalyzer (Nano RNA Chip; Agilent). Whole transcriptome libraries were prepared with the Ion Torrent Total RNA-Seq Kit v2 (Thermo Fisher Scientific) from 200ng total poly-A enriched RNA (Dynabead mRNA direct Micro Kit; Ambion). Sequencing was done on an Ion P1 chip using the Thermo Fisher Ion Proton System to a minimum of 30M reads.

#### Isolating public and superpublic CDR3s

For each cohort, CDR3 beta amino acid sequences were pooled together and occurences in the cohort of each individual sequence was counted. Public CDR3aa are defined as seen in at least two individuals in the cohort, while superpublic CDR3aa are defined as seen in at least half of the individuals in the cohort.

#### Calculating public fraction by frequency and by sequence

For each individual, the public fraction (Figure 2I) is calculated as the fraction of the CDR3aa that overlaps with the cohort’s public CDR3aa pool. The summed clonality (Figure 2J) is calculated by summing the clonality (relative CDR3 frequency in the sample) of the CDR3aa overlapping with the cohort’s public CDR3aa pool.

#### Disease-specific CDR3 sets

A set of 160,000 unique SARS-CoV2-specific CDR3 was obtained from the ImmuneCODE™ database (Nolan et al., 2020). From the *McPAS* CDR3 datasets, we downloaded the McPAS database on 2021-08-12 (Tickotsky et al., 2017). We included in our study CDR3beta amino acid sequences of human origin found in the two top categories of diseases: Pathogens and Autoimmune.

#### Isolating CDR3 from bulk RNA-Seq *in silico*

From each RNA-Seq donor sample from the *GVHD* cohort, we isolated CDR3 contigs using the MIXCR software (Bolotin et al., 2015). Since the Ion Proton sequencing system generates variable-length reads, we allowed for partial alignments and performed two passes of contig assembly. To rescue as many CDR3s as possible, for incomplete TCR CDR3s, we allowed for extension via the V/J genes, since it has been shown to introduce limited errors, because of the very conserved nature of TCRs on both ends (pattern CASS-------EF) (Bolotin et al., 2015).

#### Peptide sets and TCR-binding prediction

We downloaded 25,270 human-derived HLA-associated peptides from (Pearson et al., 2016) on 2020-06-07 and 20,961 viral-derived peptides from the Immune Epitope Database (Vita et al., 2019) on 2020-11-23. Using the ERGO model commit version 5c6fc37, cloned from the GitHub repository (https://github.com/louzounlab/ERGO), we applied the long short-term memory model pre-trained on McPAS on selected CDR3. ERGO outputs a probability of TCR recognition score between 0 and 1, where 1 means recognized.

#### Assessing polyreactivity from the tetramer study

Cells were grouped by CDR3aa sequence and peptide binding was counted with a threshold of above 0. Thus for each CDR3aa sequence and each peptide pair, all cell barcodes combined, we obtained a 0/1 value, indicating a recognition of that peptide by a cell with the CDR3aa sequence in question. This way, for each CDR3aa, we obtained a value between 0 and 50, corresponding to the 50 peptides from the *10xgenomics* study.

#### CDR3 sharing and repertoire overlaps

We calculated sharing of individual CDR3 sequences based solely on the amino acid sequence without matching V/J genes and nucleotide sequences. This approach was selected to assess sharing of the final protein product of CDR3s found in the body rather than to look at specific mRNA features.

Thus, for public fraction calculations based on unique CDR3aa sequences, we calculated the number of public sequences in an individual’s repertoire and divided it by the total number of sequences in the repertoire (results of Figure 2I). To assess the clonality of public CDR3aa, we summed the clonal frequencies attributed to individual public sequences (results of Figure 2J).

We use the Jaccard distance *d*_*j*(*A,B*_) as a measure of dissimilarity between two CDR3 sets *A* and

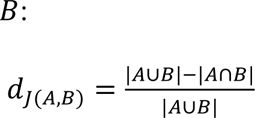

Here a value of 0 means exact overlap of the CDR3 sets, and 1 means no overlap.

#### Diversity measurements

We used the inverse Simpson clonality and Shannon entropy as diversity measurements. Inverse Simpson diversity index is calculated by weighted arithmetic mean of each squared clone abundance (Simpson, 1949), and we implemented this as a function in python. Shannon entropy (Shannon, 1948) is calculated using the *scipy.stats* python package.

#### Recombination probability prediction

CDR3aa recombination probability was predicted using the OLGA software (Sethna et al., 2019). We downloaded the command-line tool commit version 4e0bc36 from the repository (https://github.com/statbiophys/OLGA) and selected *humanTRB* as the alignment database for the predictions.

#### Number of mismatches

We used IgBLAST to align each nucleotide sequence to the annotated germline V, D, and J sequences to calculate the number of mismatches. We downloaded the package from the repository (https://github.com/ncbi/igblast) commit version dfb98f8. We used the human database and specified TCRs. We obtained the aligned sequence for each result and counted the number of mismatches between the aligned sequence and the germline.

#### Gaussian mixture model

We used the gaussian mixture model from the python *GaussianMixture* function from the *sklearn* library to fit each Gaussian Mixture Model. We selected a 2 component Gaussian Mixture Model with a diagonal covariance type. We grouped individuals of the Britanova cohort by age group. For each age group, we fitted a separate Gaussian Mixture Model on the recombination frequency and clonality quantifications. Then, we evaluated the fit of this model in other age groups. This fit was calculated as a log-likelihood of fit for the data to the pre-trained model. We then reported the average log-likelihood for each age-group - model pair in the heatmap in Figure 2.

#### Hierarchical clustering

We used hierarchical clustering with an unweighted pair group method with arithmetic mean agglomerative function for all hierarchical clustering experiments in this study. Visual assessment was used to split each dendrogram into clusters manually. We used the clustermap function from the seaborn python library to plot heatmaps and associated dendrograms.

#### Expected cumulative frequency

For each CDR3aa sequence, we calculated the sharing percentage and grouped sequences according to the CDR3aa sharing bins. Then, we calculated for each CDR3aa the cumulative repertoire frequency by summing frequencies of all CDR3aa in each bin across all individual repertoires. The average repertoire frequency for each bin was 4.12 x 10^-6^ ± 1.2 x 10^-6^. To draw the expected cumulative frequency line, we multiplied this overall frequency by the median number of individuals of each sharing bin. We reported this value as the mean expected cumulative frequency (red dotted line on the plot).

#### Overlaps by top *N* most frequent CDR3

We ranked CDR3 in descending order of clonal frequency, and for a growing *N*, we selected the top *N* most frequent CDR3 in each repertoire. Then, we calculated the percentage overlap with the disease-specific CDR3 set. Individual percentages were grouped by age group, and for each age group, the standard deviation within the age group is shown on each line plot.

#### Treemap

We used the treemap function from the *squarify* python library. The package was given the CDR3 clonal frequencies and a random color palette.

#### Survival model

We used the *survival* package in R for plotting the Kaplan-Meier plots and the *lifelines* packages in python for the CoxPH model. For each Kaplan-Meier plot, we split the group by median as well as 25% and 75% quantiles to attempt to find the best group separation for each CDR3 characteristic. All plots can be seen in Figure S6 with associated statistical testing.

For the *CoxPH* models, we used the *lifelines* python library with the option for right-censored data. On each plot, we reported the log (hazard ratios) as well as the bottom and top 95% confidence intervals.

#### Classification and regression models

The logistic regression and random forest models from the *sklearn* python library were used to classify *neonatal* and *TDT-dependent* CDR3s in Figure 7. We used the default parameters for each model: respectively, an L2 penalty with a regularization strength of 1 and the L-BFGS solver for the logistic regression and 100 estimators, a Gini impurity criterion, no max depth, and a minimum number of samples of 2 for the split for the random forest classifier.

For the ablation study, each model received the selected combination of features and learned to classify CDR3 into two classes: *neonatal* or *TDT-dependent*. During the dataset preparation, two repertories of each type (cord blood and child) were held out. These repertoires comprised the test set of new data. The performance of each iteration of the model given the combination of input features was reported in Figure 7, with performances color-coded for visual comparison.

A linear regression model was trained to determine the relative importance of the features. We used the *LinearRegression* function from the *sklearn* python library with the following default features: intercept fitting was allowed, and negative coefficients were allowed. This model received as input a binary vector of the presence/absence of the features and learned to predict the performance of either of the two models (logistic regression or random forest) obtained previously for each feature combination. The coefficients attributed to each binary feature presence/absence were used to compare relative importance. Coefficients close to zero meant there was little weight attributed to the model and vice versa. The ranking of features was obtained by ordering the absolute values of the coefficients in descending order.

### Quantification and Statistical Analysis

All statistical analyses were performed using Python v3.7.6 or R v4.0.4. All statistical tests used are mentioned in the figure legends. Significance level (p<0.05) results are marked with (*) in the figures. Mann-Whitney U and One-way ANOVA tests were performed using the mannwhitneyu and anova functions respectively from scipy.stats python module and R. All boxes in boxplots show the first (25^th^ percentile) and third quartiles (75^th^ percentile) and the median while the whiskers designate the minimum (first quartile value – 1.5*interquartile range) and maximum (third quartile value + 1.5*interquartile range).

