## Supplemental Materials for "Two types of human TCR differentially regulate reactivity to self and non-self antigens"

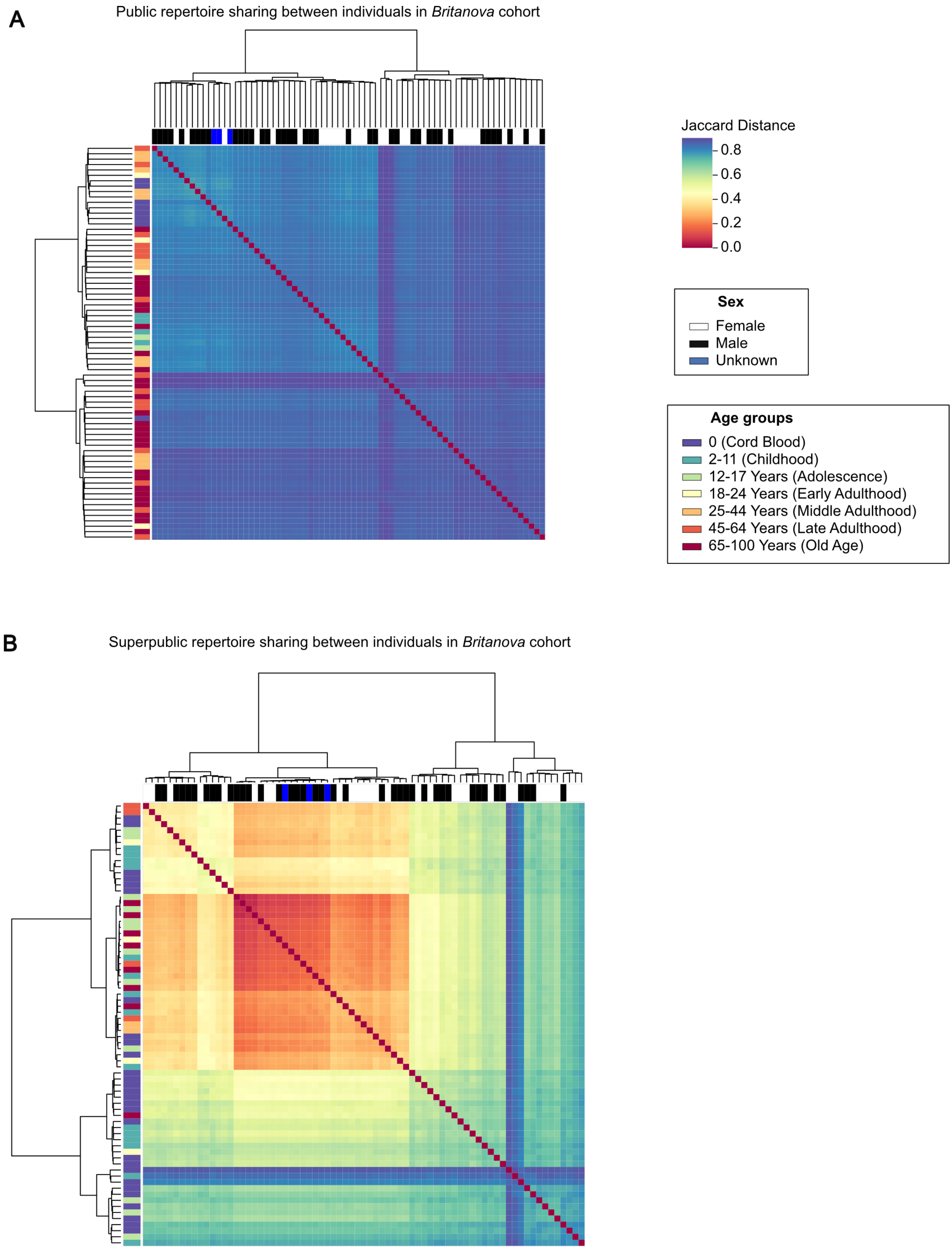


**Figure S1**: Hierarchical clustering of repertoire sharing (pairwise Jaccard distance) among subjects of the Britanova cohort for **(A)** public and **(B)** superpublic CDR3s.

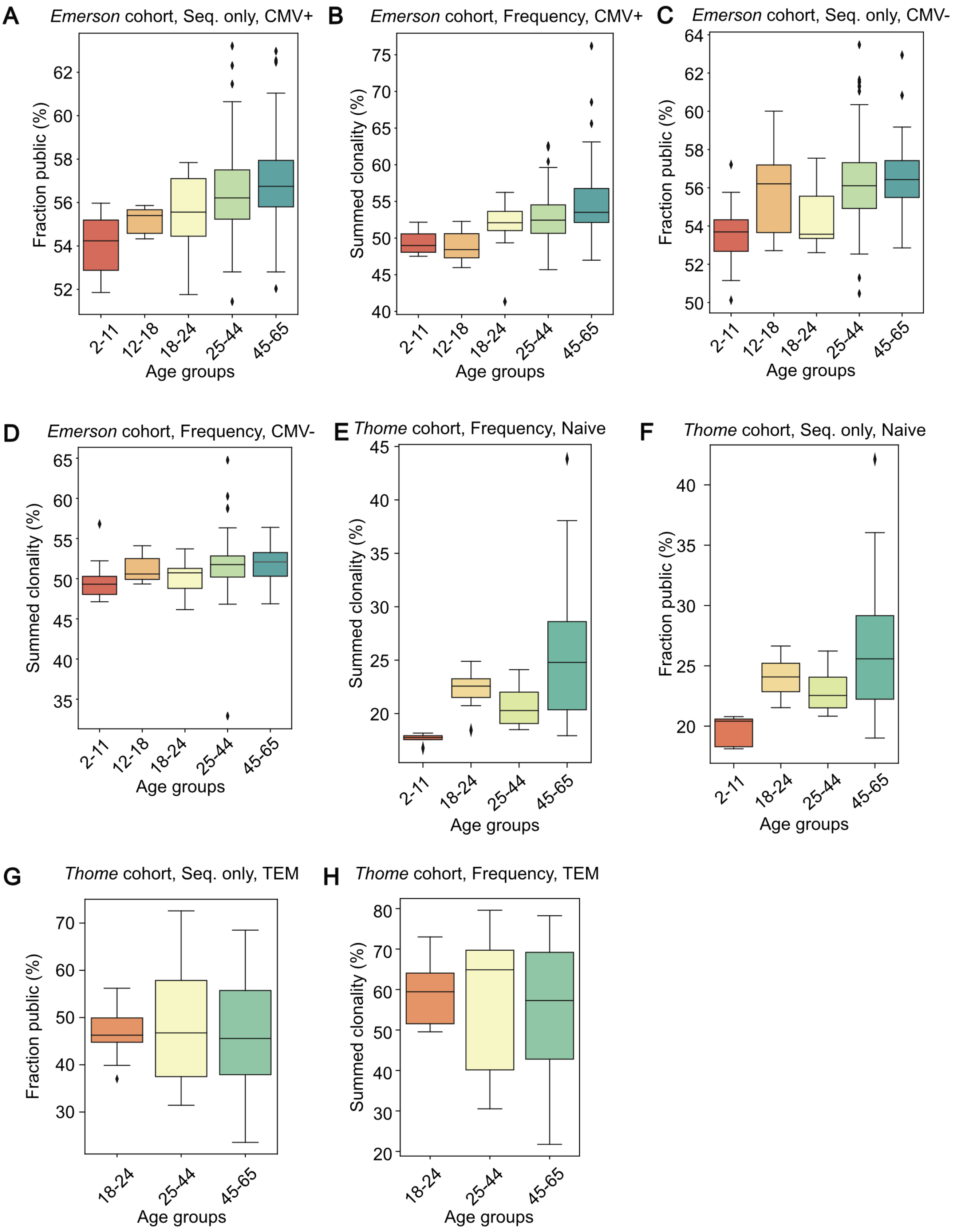


**Figure S2**: Boxplots showing **(A-C)** public fraction percentages and **(B-D)** summed clonality of public CDR3aas in the Emerson cohort CMV+ and CMV- individuals. Boxplots showing **(E-G)** public fraction percentages and **(F-H)** summed clonality of public CDR3aas for naïve and effector-memory T cells in the *Thome* cohort.

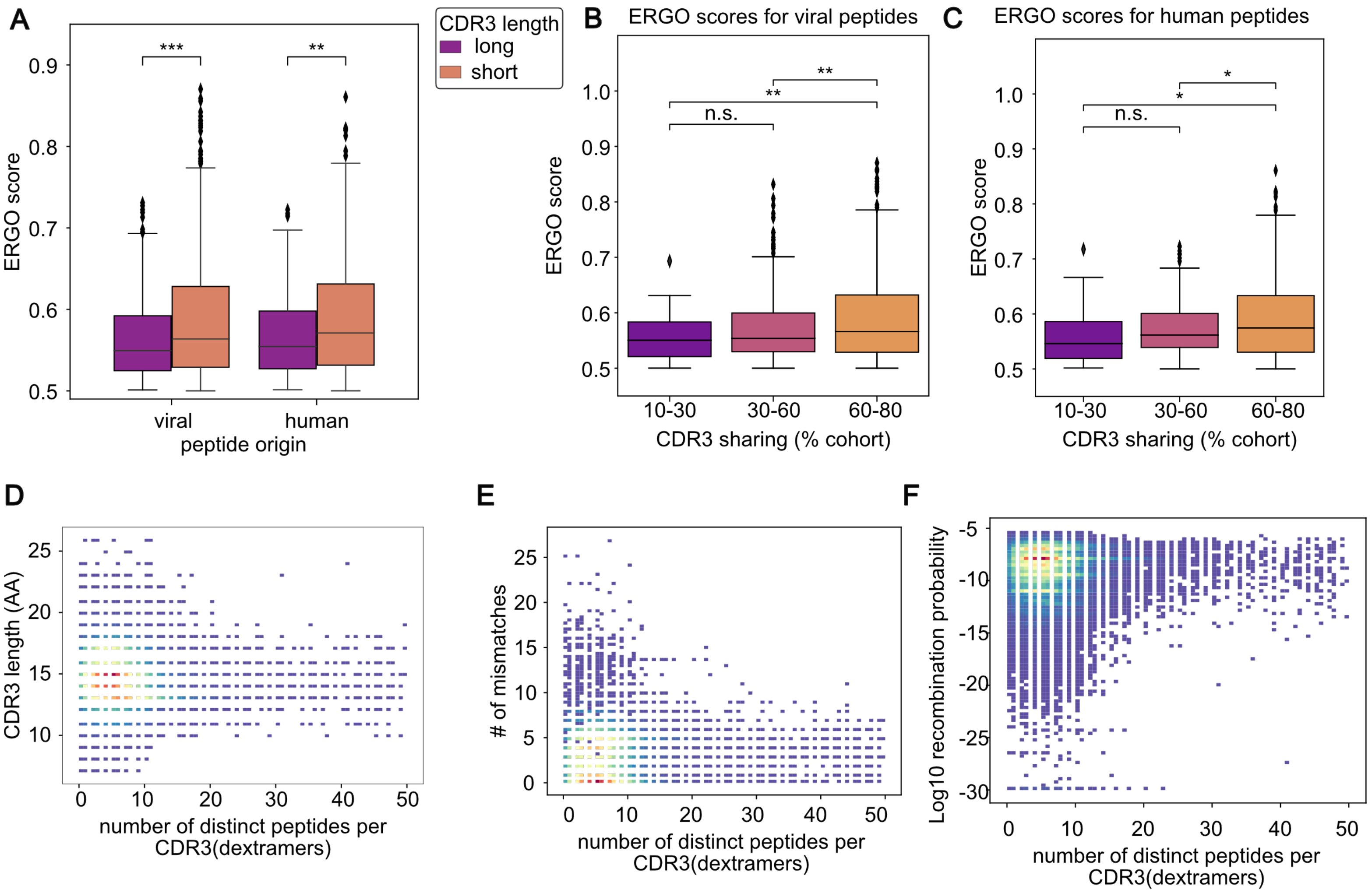


G

*Britanova* cohort broad age groups vs sex

| | Male | Female | Unknown |
|---|---|---|---|
| 0 (Cord blood) | 3 | 2 | 3 |
| 2-44 (Young) | 12 | 10 | 0 |
| 45-64 (Adult) | 5 | 10 | 0 |
| 65+ (Old Age) | 10 | 18 | 0 |

*Comparison of CDR3 repertoire size between cohorts*

| | Mean # CDR3aa |
|---|---|
| Britanova | 339383±233324 |
| Emerson | 198819±83149 |
| Thome | 19645±14952 |
| Sng | 20597±12676 |

**Figure S3**: **(A)** ERGO scores were used to predict the binding of CDR3aa longer or shorter than 15 amino acids to MHC-associated peptides of viral and human origin. ERGO scores were used to assess the relationship between two features of CDR3aa from the Britanova cohort: publicness and polyreactivity to **(B)** viral and **(C)** human peptides. Polyreactivity vs. **(D)** CDR3aa length, **(E)** the number of mismatches, and **(F)** log10 recombination frequency in the 10x Genomics dataset. **(G)** Distribution of *Britanova* cohort individuals by sex and with broad age groups depicted in Figure 3. **(H)** Comparison of CDR3aa repertoire sizes by cohort.

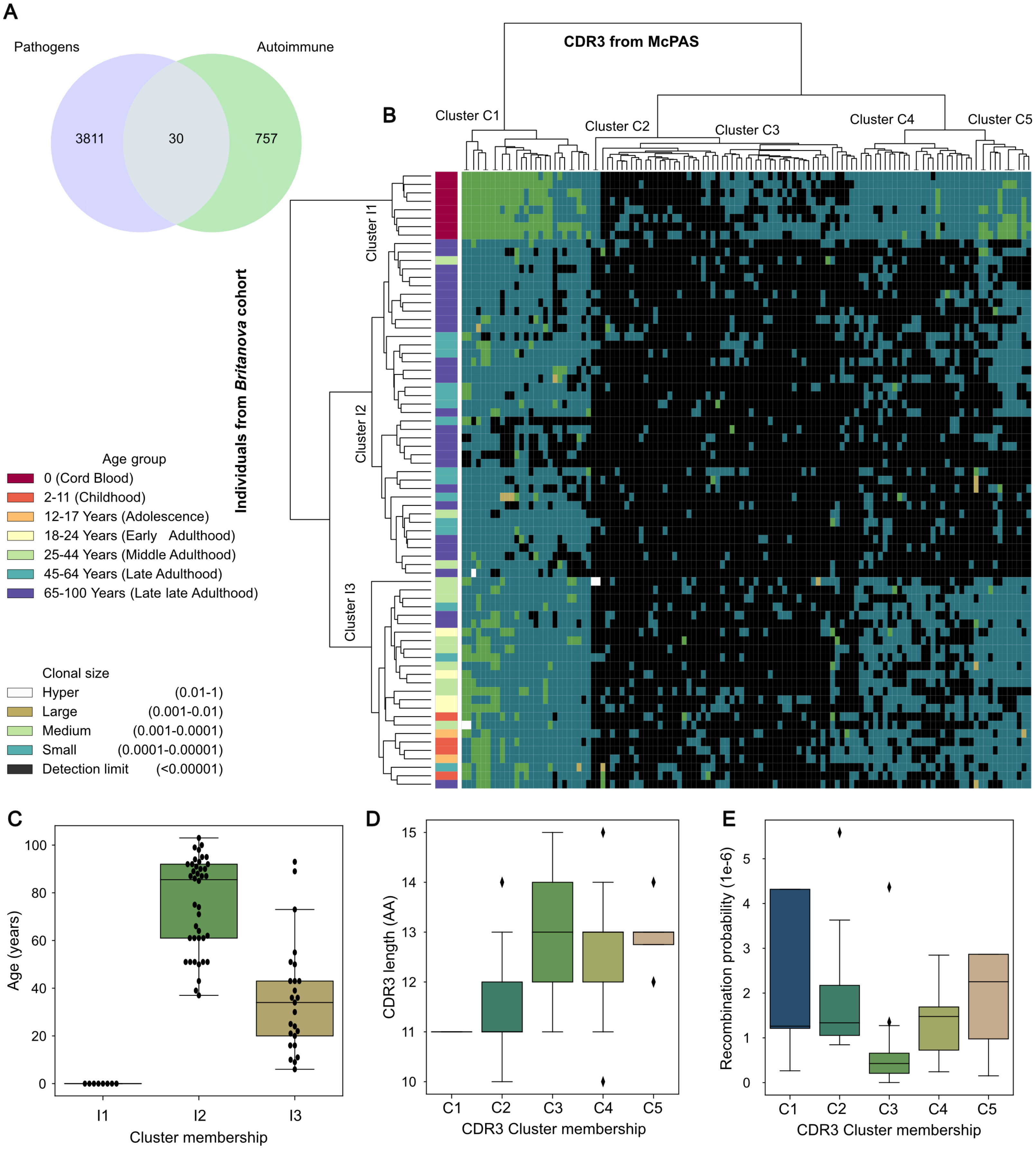


**Figure S4**: **(A)** Venn diagram showing the overlap between two major CDR3 categories from McPAS: autoimmunity and microbial pathogens. **(B)** Heatmap shows, for subjects of the Britanova cohort, the frequency of CDR3aa listed in the McPAS autoimmune dataset. Rows represent individuals, columns unique CDR3aa, and cell color indicates CDR3aa clone size. Row dendrogram leaves are colored by age group. **(C)** Age distribution for subjects in three individual clusters from **(A)**. **(D)** Boxplot showing CDR3aa length in five clusters from **(B)**. **(E)** Boxplot showing predicted recombination frequency for CDR3aa in five clusters from **(B)**.

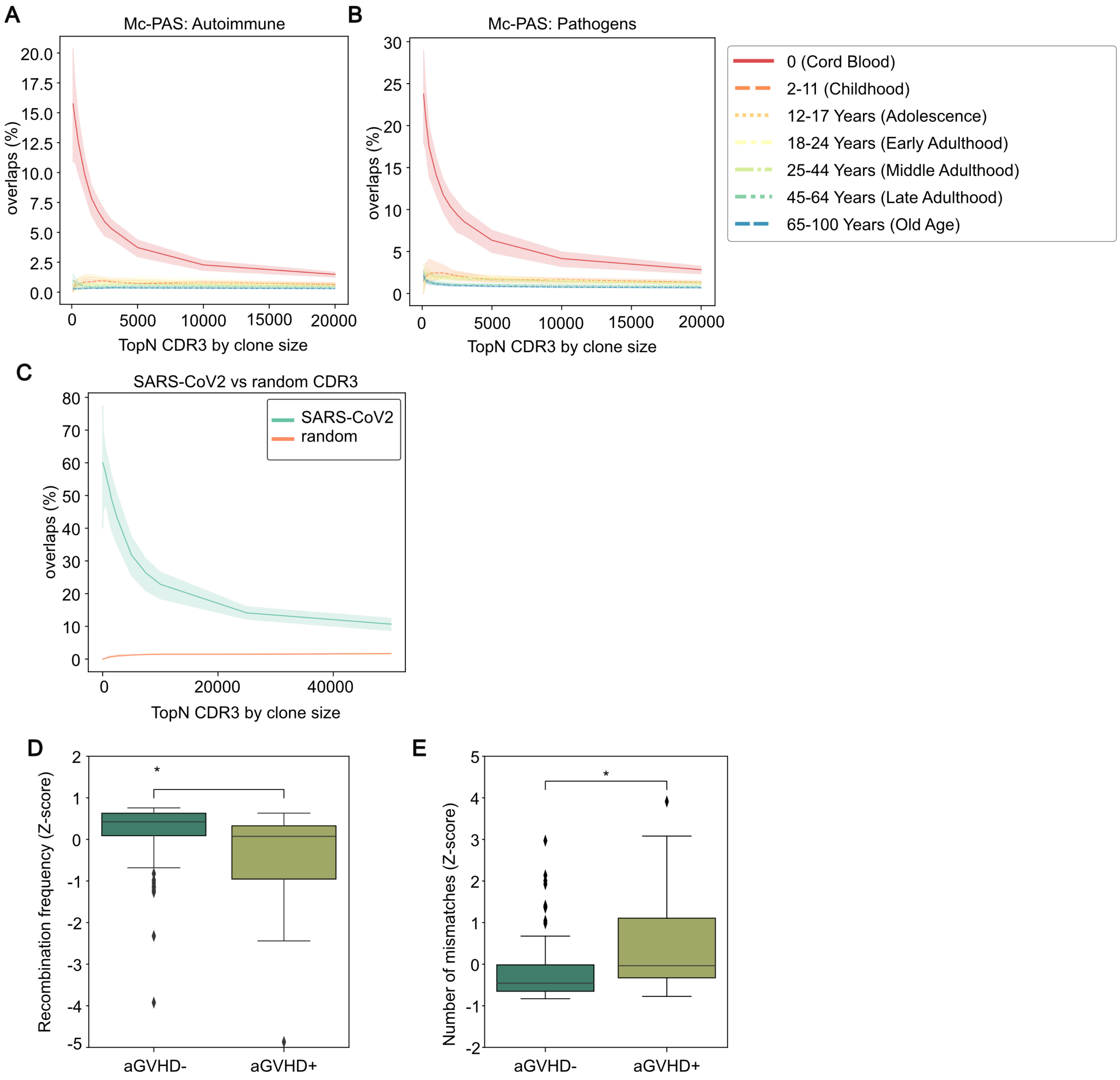


**Figure S5**: Line plots showing percentage overlaps with McPAS **(A)** Pathogen-specific and **(B)** Autoimmune-specific CDR3 set, with the top N most frequent CDR3aa. Line colors and types correspond to age groups. **(C)** SARS-CoV2-specific CDR3 overlaps vs. a randomly selected set of CDR3 in cord blood. Boxplots showing **(D)** the recombination frequency and **(E)** the number of mismatches for CDR3aa in aGVHD+ vs. aGVHD- donors.

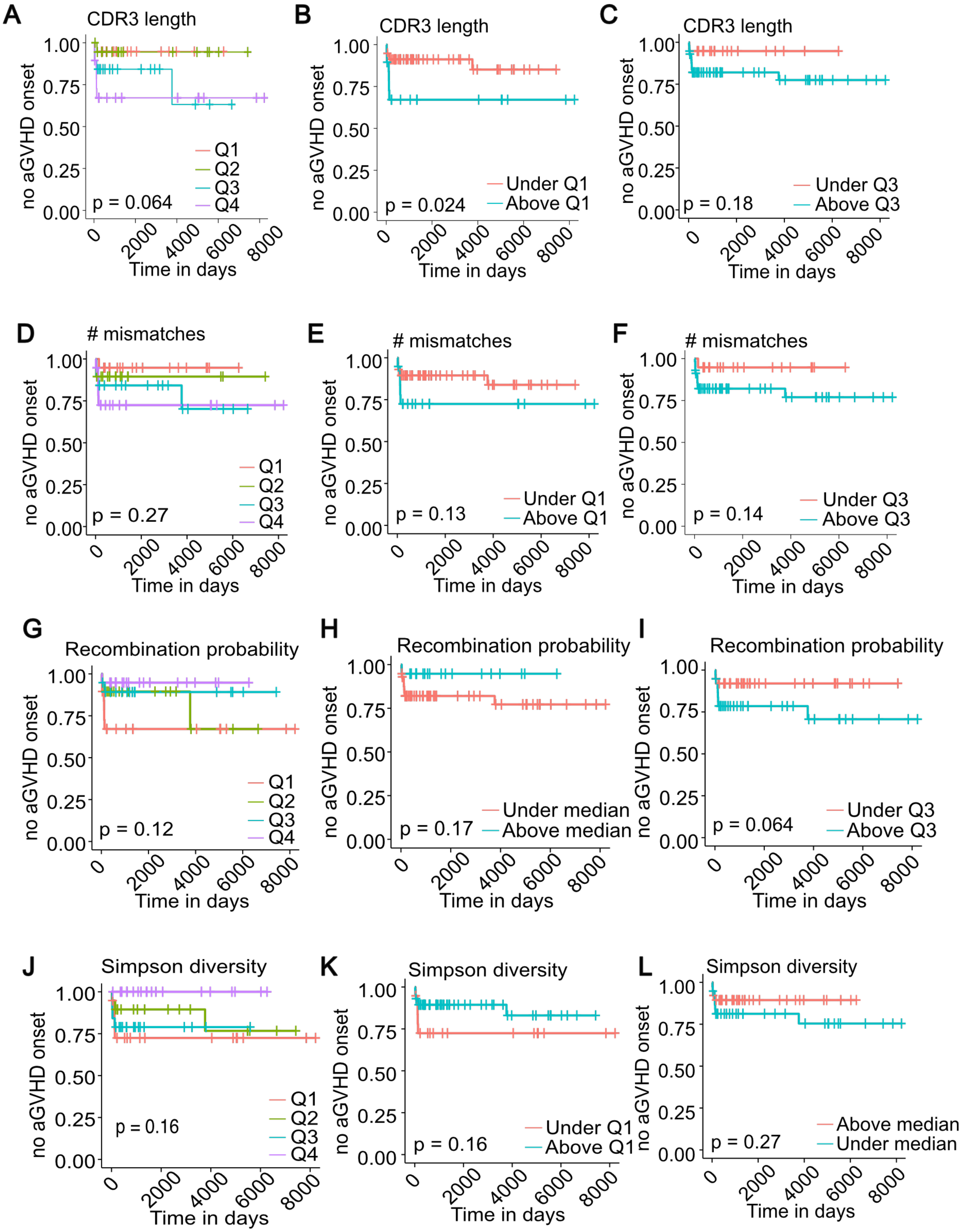


**Figure S6**: Kaplan-Meier plots showing splits by all four and individual quartiles for **(A-C)** CDR3 length, **(D-F)** the number of mismatches, **(G-I)** recombination probability, and **(J-L)** Simpson diversity. This figure complements the results from Figure 6.

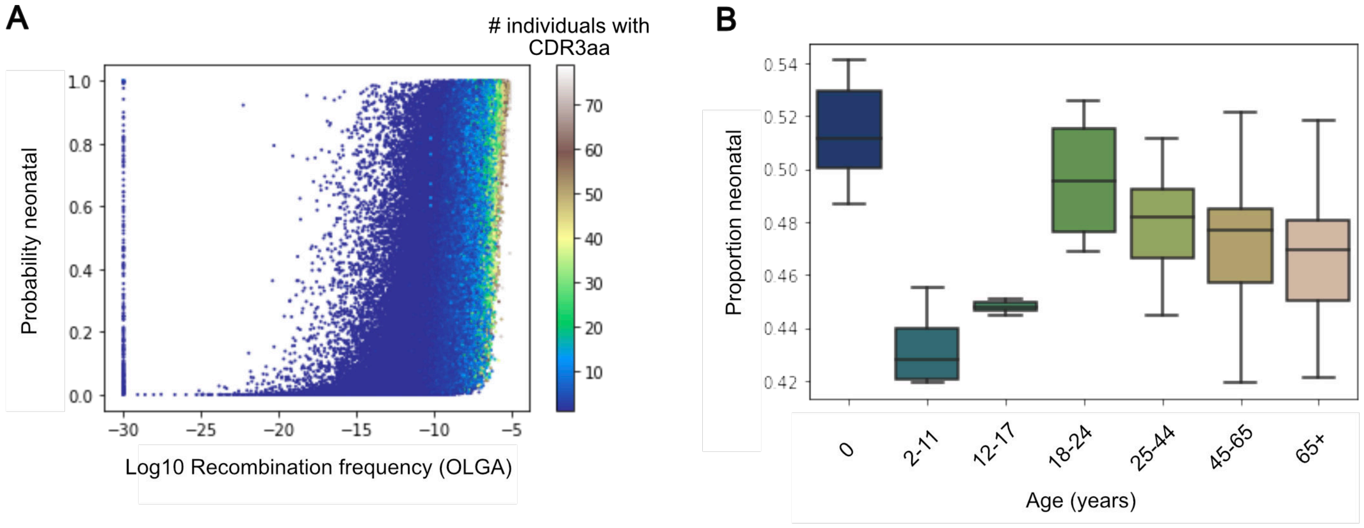


**Figure S7**: **(A)** Scatterplot showing the recombination frequency for CDR3s based on the classification probability output of the logistic regression model (neonatal vs. TDT-dependent). Each dot is a CDR3s, and the color scheme represents the degree of sharing through the cohort. **(B)** The proportion of neonatal CDR3s by age group in the Britanova cohort.
